# Neuronal modeling of magnetoencephalography responses in auditory cortex to auditory and visual stimuli

**DOI:** 10.1101/2023.06.16.545371

**Authors:** Kaisu Lankinen, Jyrki Ahveninen, Mainak Jas, Tommi Raij, Seppo P. Ahlfors

**Affiliations:** Athinoula A. Martinos Center for Biomedical Imaging, Massachusetts General Hospital, Charlestown, MA 02129; Department of Radiology, Harvard Medical School, Boston, MA 02115

**Author notes:** Corresponding author: Kaisu Lankinen.

## Abstract

Previous studies have demonstrated that auditory cortex activity can be influenced by crosssensory visual inputs. Intracortical recordings in non-human primates (NHP) have suggested a bottom-up feedforward (FF) type laminar profile for auditory evoked but top-down feedback (FB) type for cross-sensory visual evoked activity in the auditory cortex. To test whether this principle applies also to humans, we analyzed magnetoencephalography (MEG) responses from eight human subjects (six females) evoked by simple auditory or visual stimuli. In the estimated MEG source waveforms for auditory cortex region of interest, auditory evoked responses showed peaks at 37 and 90 ms and cross-sensory visual responses at 125 ms. The inputs to the auditory cortex were then modeled through FF and FB type connections targeting different cortical layers using the Human Neocortical Neurosolver (HNN), which consists of a neocortical circuit model linking the cellular– and circuit-level mechanisms to MEG. The HNN models suggested that the measured auditory response could be explained by an FF input followed by an FB input, and the crosssensory visual response by an FB input. Thus, the combined MEG and HNN results support the hypothesis that cross-sensory visual input in the auditory cortex is of FB type. The results also illustrate how the dynamic patterns of the estimated MEG/EEG source activity can provide information about the characteristics of the input into a cortical area in terms of the hierarchical organization among areas.

**Significance statement:** Laminar intracortical profiles of activity characterize feedforward– and feedback-type influences in the inputs to a cortical area. By combining magnetoencephalography (MEG) and biophysical computational neural modeling, we obtained evidence of cross-sensory visual evoked activity in human auditory cortex being of feedback type. The finding is consistent with previous intracortical recordings in non-human primates. The results illustrate how patterns of MEG source activity can be interpreted in the context of the hierarchical organization among cortical areas.

## Introduction

Activity in sensory cortices is influenced by feedforward (FF) and feedback (FB) connections between cortical layers and brain regions, following a hierarchical organization (Rockland and Pandya, 1979; Felleman and Van Essen, 1991; Zeki, 2018). In the auditory cortex of non-human primates (NHPs), the laminar profile of early auditory evoked responses has FF type characteristics, whereas cross-sensory visual or somatosensory evoked activity are of FB type (for reviews see, e.g., Foxe and Schroeder, 2005; Schroeder and Foxe, 2005; Ghazanfar and Schroeder, 2006; Kayser and Logothetis, 2007). Human magneto– and electroencephalography (MEG/EEG) studies have revealed that cross-sensory activations and multisensory interactions can occur in low-order sensory areas very early, within a few tens of milliseconds from the stimulus onset (Giard and Peronnet, 1999; Foxe et al., 2000; Molholm et al., 2002; Teder-Sälejärvi et al., 2002; Molholm et al., 2004; Lakatos et al., 2007; Talsma et al., 2007; Raij et al., 2010). In line with evidence from studies in other cognitive domains (Polimeni et al., 2010; Muckli et al., 2015; Kok et al., 2016; Fracasso et al., 2018; Klein et al., 2018; Finn et al., 2019; Lawrence et al., 2019a; Norris and Polimeni, 2019), recent high-field fMRI studies have provided evidence of FF– and FB-like intracortical depth profiles in auditory cortex BOLD signals (De Martino et al., 2015; Ahveninen et al., 2016; Moerel et al., 2018; Wu et al., 2018; Moerel et al., 2019; Gau et al., 2020; Chai et al., 2021; Lankinen et al., 2022). However, detailed neurophysiological analysis or computational modeling of such effects has not been done in humans.

Previous studies have suggested that early components of evoked responses are related to FF processes, whereas later components reflect FB influences in activity evoked by auditory (Inui et al., 2006; Kohl et al., 2022), visual (Aine et al., 2003; Inui and Kakigi, 2006), and somatosensory (Cauller and Kulics, 1991; Inui et al., 2004; Jones et al., 2007) stimuli.

Biophysically realistic computational models have been used to investigate laminar connections and cellular and circuit level processes of the neurons in detail, and they can also be used to simulate MEG/EEG signals (Jones et al., 2007; Neymotin et al., 2020). The Human Neocortical Neurosolver (HNN) (Neymotin et al., 2020) provides a cortical column model with FF– and FB-type inputs targeting different layers. With HNN, the cellular and network contributions to MEG/EEG signals from a source-localized region of interest can be modeled and compared to the measured signals. Previously, HNN has been used to interpret mechanisms of sensory evoked responses and oscillations in healthy and clinical populations (Jones et al., 2007; Jones et al., 2009; Ziegler et al., 2010; Lee and Jones, 2013; Khan et al., 2015; Sherman et al., 2016; Pinotsis et al., 2017; Sliva et al., 2018; Bonaiuto et al., 2021; Kohl et al., 2022; Law et al., 2022). Kohl et al. (2022) showed that auditory responses in the auditory cortex could be modeled by activating the neocortical circuit through a layer-specific sequence of FF-FB-FF inputs, similar to a prior simulation of somatosensory evoked responses (Jones et al., 2007).

In the present study, we investigated auditory vs. cross-sensory visual evoked responses in the auditory cortex by comparing the measured MEG responses with simulated source waveforms from a computational model (HNN). We hypothesized that the auditory evoked responses observed with MEG can be explained by a sequence of FF and FB influences, whereas FB-type input is adequate to explain the cross-sensory visual evoked response.

## Material and methods

### Subjects

Eight healthy right-handed subjects participated (six females, age 22–30 years). All subjects gave written informed consent, and the study protocol was approved by the Massachusetts General Hospital institutional review board and followed the guidelines of the Declaration of Helsinki.

### Stimuli and task

The subjects were presented with *Noise/Checkerboard* and *Letter* stimuli in separate runs while MEG was recorded. Data for the *Noise/Checkerboard* stimuli were used in our earlier publication (Raij et al., 2010). Here we re-analyzed data from the *Noise/Checkerboard* experiment, together with the previously unpublished data from the *Letter* experiment. Equiprobable 300-ms auditory, visual, and audiovisual (simultaneous auditory and visual) stimuli were delivered in an eventrelated design with pseudorandom order. The auditory *Noise* stimuli were white noise bursts (15 ms rise and decay) and the visual *Checkerboard* stimuli static checkerboard patterns (visual angle 3.5°×3.5° and contrast 100%, with a peripheral fixation crosshair). The *Letter* stimuli were spoken and written letters of Roman alphabet (‘A’, ‘B’, ‘C’, etc.). The subjects’ task was to respond to rare (10%) auditory, visual, or audiovisual target stimuli with the right index finger movement as quickly as possible. In the *Noise/Checkerboard* experiment, the target stimulus was a tone pip, a checkerboard with a gray diamond pattern in the middle, or a combination of the two. In the *Letter* task, the target stimulus was the letter ‘K’, spoken and/or written. Data were recorded in three runs with different stimulus onset asynchrony (SOA, mean 1.5, 3.1, or 6.1 s, all jittered at 1.15 s). There were 375 stimuli per category (auditory, visual, and audiovisual): 150 in the short, 125 in the intermediate, and 100 in the long SOA runs. All subjects were presented with the same order of tasks and stimuli. The auditory stimuli were presented with MEG-compatible headphones, with the intensity adjusted to be as high as the subject could comfortably listen to. The visual stimuli were projected onto a translucent screen. The stimuli were controlled using Presentation 9.20 (Neurobehavioral Systems Inc, Albany, CA, USA).

### MEG and MRI acquisition and co-registration

MEG was recorded with a 306-channel instrument with 204 planar gradiometer and 102 magnetometer sensors (VectorView; MEGIN, Finland) inside a magnetically shielded room (Cohen et al., 2002). Simultaneous horizontal and vertical electro-oculograms (EOG) were also recorded. All signals were bandpass-filtered to 0.03–200 Hz and sampled at 600 Hz.

Structural T1-weighted MRIs of the subjects were acquired with a 1.5 T Siemens Avanto scanner (Siemens Medical Solutions, Erlangen, Germany) and a head coil using a standard MPRAGE sequence. Cortical surfaces were reconstructed using the FreeSurfer software (http://www.surfer.nmr.mgh.harvard.edu, (Fischl, 2012).

Prior to the MEG recording, the locations of four small head position indicator coils attached to the scalp and several additional scalp surface points were determined with respect to the fiducial landmarks (nasion and two preauricular points) using a 3-D digitizer (Fastrak Polhemus, VT, USA). For the MRI–MEG coordinate system alignment, the fiduciary points were first identified from the structural MRIs, and then this initial co-registration was refined using an iterative closestpoint search algorithm for the scalp surface locations using the MNE Suite software (Gramfort et al., 2014, http://www.martinos.org/mne/).

### MEG preprocessing and source estimation

The MEG data were analyzed using MNE-Python (Gramfort et al., 2013). After excluding channels and time segments with excessive noise, independent component analysis (ICA) was used to identify and remove artifacts related to eye blinks, eye movements, and cardiac activity. The signals were then lowpass filtered at 40 Hz, and event-related responses were averaged separately for the auditory and visual trials, combining the long, intermediate, and short SOA runs.

After exclusion of artifactual time segments an average of 369.9 (std 6.5) epochs per subject remained in response to auditory, and 370.2 (std 5.1) to visual stimulation. In the present study we did not analyze the audiovisual or target trials. The zero level in each channel was defined as the mean signal over the 200-ms prestimulus baseline period.

Source activity was estimated at 4098 discrete locations per hemisphere on the cortical surface, with an average separation of the source elements being about 4.9 mm. For the forward solution, a single-compartment boundary element model was used. Forward solutions were first computed separately for the three runs with different SOAs and then averaged (Uutela et al., 2001). Minimum-norm estimates (MNE, (Hamalainen and Ilmoniemi, 1994)) for the cortical source currents were calculated. Both the gradiometer and the magnetometer channels were included in the source estimation. We used fixed source orientation normal to the cortical surface and depth weighting 0.8 to reduce bias towards superficial currents. For region-of-interest (ROI) selection, the MNE values were noise-normalized to obtain dynamic statistical parametric maps (dSPM; Dale et al., 2000).

### Regions-of-interest and source time courses

Auditory evoked potentials and magnetic fields typically have three main deflections: P50-N100P200 (or P50m-N100m-P200m for MEG), peaking approximately at 50, 100 and 180 ms, respectively, after the auditory stimulus onset (Picton et al., 1974; Hari et al., 1980; Hämäläinen et al., 1993; Jones et al., 2007; Ahlfors et al., 2015). The ROIs were determined based on the auditory N100m response, because the SNR of the visual evoked response over the auditory cortex was too low to reliably determine auditory cortex ROIs from the visual evoked data in the presence of partially coinciding strong occipital visual cortex activity. We identified functional ROIs for the auditory cortex in each hemisphere, separately for each subject, based on the N100m peak of the auditory evoked response. First, anatomically defined regions were selected using the Destrieux atlas parcellation from Freesurfer (Fischl et al., 2004; Destrieux et al., 2010):

Heschl’s gyrus, Heschl’s sulcus, and the lower part of planum temporale (masked with supramarginal gyrus) were combined to cover the primary auditory areas. Then, from these regions the source element with the largest negative deflection between 60–110 ms (except for manually set 105 ms in one subject) in the dSPM source time course was identified. Using that source element as a seed point, all source elements that had a magnitude of 30% or more of the peak dSPM value and formed a continuous area around the seed point were selected. The average number of selected elements across subjects, hemispheres and experiments for the auditory cortex ROIs was 19 (standard deviation 8.7, range 3–38). The same procedure was used to determine also additional control ROIs in the occipital cortex (V1, V2, and MT based on the FreeSurfer atlas (Fischl et al., 2008). The source waveform for an ROI was defined as the sum of the MNE time courses over those selected source elements. Note that the magnitude of the response depended on the number of the vertices that were included in the ROI, and thus was expected to give a smaller amplitude than would be found by the use of a single equivalent current dipole to represent the auditory cortex activity (as used, *e.g.*, by Kohl et al. (2022)). Although equivalent current dipoles are in general well suited to describe auditory evoked responses, here it was more convenient to use a distributed source model (MNE) for wide-spread visual evoked response, to extract cross-sensory responses in the auditory cortex.

### Neural modeling with Human Neocortical Neurosolver (HNN)

Activity in the auditory cortex evoked by the auditory and visual stimuli was modeled using HNN (https://jonescompneurolab.github.io/hnn-core/) (Neymotin et al., 2020). HNN is a software for simulating neocortical circuits and linking cellular– and circuit-level physiology to the electrical source currents measured by MEG and EEG. Thus, HNN provides a tool to develop and test hypotheses on the neural origins of MEG/EEG signals. The neural currents contributing to the MEG/EEG signals from a source region are modeled in terms of the local network dynamics driven by layer-specific inputs (see **Fig. 1**). Simulated MEG/EEG source currents are represented as current dipole waveforms calculated from the distribution of intracellular currents in the dendrites of the pyramidal cells. MEG/EEG signals originate mostly from postsynaptic currents in cortical pyramidal neurons (Hämäläinen et al., 1993; Okada et al., 1997), and the magnitude and direction 193 of the source current depends on the type of the synaptic input and its dendritic location (Allison 194 et al., 2002; Jones et al., 2007; Linden et al., 2010; Lopes da Silva, 2010; Ahlfors et al., 2015; Ahlfors and Wreh, 2015), providing a link between the laminar distribution of synaptic inputs and he MEG/EEG source waveforms.

**Figure 1.**
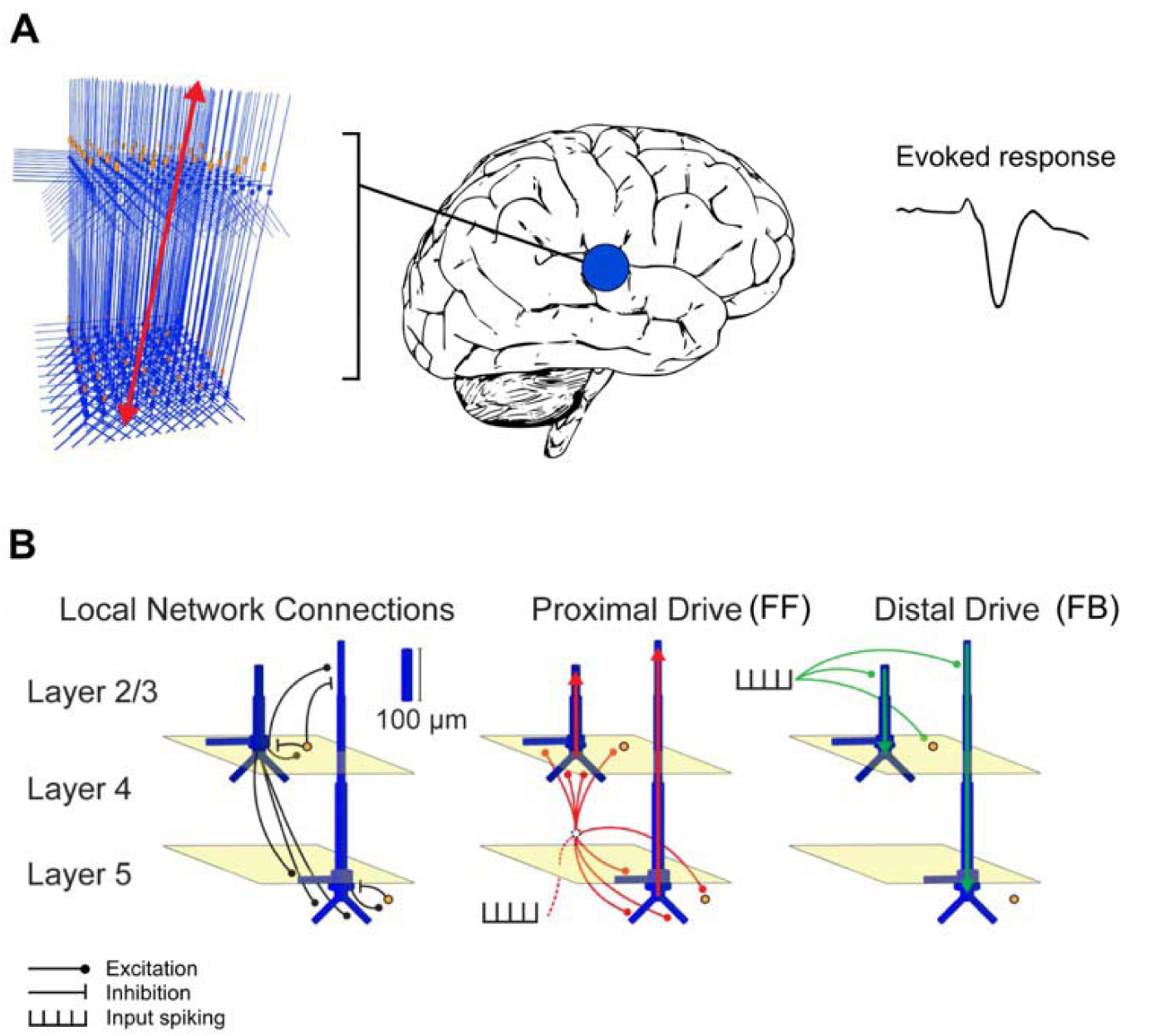
Schematic illustration of the HNN model. (A) A network of neurons in a local cortical area generates an evoked response. (B) Local network structure with pyramidal cells (blue) and 200 interneurons (orange). Excitatory and inhibitory coupling is indicated by a black circle and bar, 201 respectively. The network is activated by proximal (red) and distal (green) drives by input spike trains. Modified from Neymotin et al. (2020).

In HNN, the model for a local cortical circuit has a layered structure with pyramidal neurons whose somata are in the supragranular (layer 2/3) or infragranular (layer 5) layers and whose dendrites span across the layers. The model also includes inhibitory interneurons. External input to the circuit arrives through characteristic layer-specific FF and FB type connections. FF type inputs consist of proximal drives to the basal dendrites of the pyramidal cells (assumed to arrive via the middle cortical layer), whereas FB inputs are represented by distal drive to the apical dendrites of the pyramidal cells. The model has 100 pyramidal neurons in each of layers 2/3 and 5; a scaling factor is used to match the simulated dipole to the magnitude of the recorded evoked response. The parameters of the HNN model originate from known anatomical and physiological cell properties, and the local connectivity within and between cortical layers is based on a large body of literature from animal studies (Jones et al., 2007; Neymotin et al., 2020).

We used HNN to test the hypothesis that the differences in the MEG responses to auditory and visual stimuli can be explained by a different sequence of FF and FB inputs to the auditory cortex. This hypothesis is based on neurophysiological evidence from animal studies (Schroeder and Foxe, 2002). Underlying mechanisms of auditory responses in humans have been previously described using HNN (Kohl et al., 2022). Our specific hypothesis was that the auditory response can be explained by an initial FF input followed by an FB input, but the visual response just by an FB input.

We created two main HNN models for event-related activity in the auditory cortex: one for the response to auditory stimuli and one for the response to visual stimuli. The grand average MEG source waveforms (averaged across subjects, hemispheres, and experiments) were modeled using HNN. As a starting point, we used the auditory cortex model by Kohl et al. (2022) for activity in the right hemisphere evoked by auditory stimuli presented to the left ear. Because HNN has a large number of user-defined parameters, we made the following assumptions to limit the parameter space: a) Only the timing parameters of the FF/FB spike-train inputs (mean µ and standard deviation s of a Gaussian distribution) were adjusted, in addition to an overall scaling factor for the simulated source waveforms; all the other parameters were kept unchanged. b) These other, internal, model parameters were assumed to be the same for the responses to visual and auditory stimuli. c) The simulations were limited to the time window of 0–150 ms for the auditory and 0–170 ms for the visual response, in order to focus on the early part of the responses. HNN model parameters were determined by minimizing the root mean square error (RMSE) between the simulated and experimentally observed MEG source waveforms. To improve the SNR of the experimental data, we averaged MEG source waveforms over subjects, hemispheres, and the two experiments. The simulated HNN waveforms were smoothed in the default 30-ms window (Hamming window convolution).

We first manually adjusted the start time of the FF/FB inputs and scaling of the response to achieve a close initial fit to the MEG responses. An optimal scaling factor was determined by minimizing the RMSE between the average of 10 simulation runs and the MEG waveform over the specified time windows. Thereafter, we further tuned the model parameters using Bayesian optimization implemented in scikit-optimize (Head, 2020) (http://doi.org/10.5281/zenodo.1207017) for estimating µ (mean input spike timing) and σ (temporal distribution of input spikes) for each model by minimizing the RMSE between the simulated and the measured signal. We used “expected improvement” as the acquisition function.

The initial parameters were defined from the manual fit and the bounds for the search space were (µ_!!_: 20…50, µ_!“_: 55…95, µ_!!#_: 90…130, σ_!!_: 1…5, σ_!“_: 5…20, σ_!!#_: 5…20).

As HNN has a large number of parameters, it is possible that even after optimizing our main models, some other combination of parameter values could explain the waveforms equally well or better. Therefore, we formed alternative models by varying the number and timing of the FF and FB inputs. We focused on the comparison of FF + FB *vs.* FB models for explaining the early part of the MEG activity evoked by auditory and visual stimuli.

### Statistical analyses

To evaluate whether the magnitudes of the estimated MEG source waveforms (averaged across tasks and hemispheres) were significantly different from zero, we used *t*-tests with a threshold *p* < 0.05 in each of the 150 time points in the 0–250 ms window. The *p*-values were Bonferroni adjusted for the two stimulus types and 150 time points. To evaluate between-subject consistency of the magnitudes of the largest defections in the evoked responses in each hemisphere and experiment, the average value over time points within ±10 ms windows around the peak latencies were calculated for each subject and submitted to *t*-tests with False discovery rate (*fdr)* adjustment for 12 tests.

For the HNN models, a non-parametric resampling approach was used to test whether the alternative models could provide a significantly better fit than our main models. First, the MEG source waveforms for auditory and visual evoked responses were resampled by drawing from 32 signals (8 subjects x 2 hemispheres x 2 experiments) 500 times with replacement. The same was repeated for 32 simulation runs for each of the models (FF + FB and FB). Next, the root-meansquare error (RMSE) between each of the 500 resampled MEG signals and 500 resampled simulations for each model was calculated, resulting in histograms of RMSE values within each model. We tested whether the difference between the simulated source waveforms from the FB *vs.* FF + FB models was significantly different from 0. The RMSE difference histograms were normalized for each model between –1 and 1, as the ranges in the auditory and visual models were different. To create a null-distribution, the signs of the waveforms were randomly flipped 10,000 times, an average of 500 resamplings was calculated. To assign a *p*-value for each model, the mean RMSE value was compared with the null distribution, with the Bonferroni adjustment of *n* = 2 (auditory and visual models). If the difference of the models (FF + FB vs. FB) was significant, we concluded that including the first FF was necessary for the model.

## Results

### MEG source waveforms in auditory cortex in response to auditory and visual stimuli

Estimated MEG source waveforms for auditory and visual evoked activity in the auditory cortex ROIs, averaged over subjects, tasks, and hemispheres, are shown in **Fig. 2**. The auditory evoked response showed a characteristic biphasic P50m-N100m waveform, with a positive peak at 37 and a negative peak at 90 ms after the onset of the auditory stimuli. These peak latencies are similar to those reported previously for auditory noise burst stimuli (Hari et al., 1987). The crosssensory visual evoked response in the auditory cortex had a monophasic peak at 125 ms after the appearance of the visual stimuli. The source magnitudes at the peak latencies were significantly different from zero (*t*-test, *p* < 0.05, Bonferroni adjusted). The magnitude of the visual evoked response was about 13% of the magnitude of the auditory N100m. The direction of the source current for the visual response was the same as that of the auditory N100m response, pointing from the gray matter towards the white matter.

**Figure 2.**
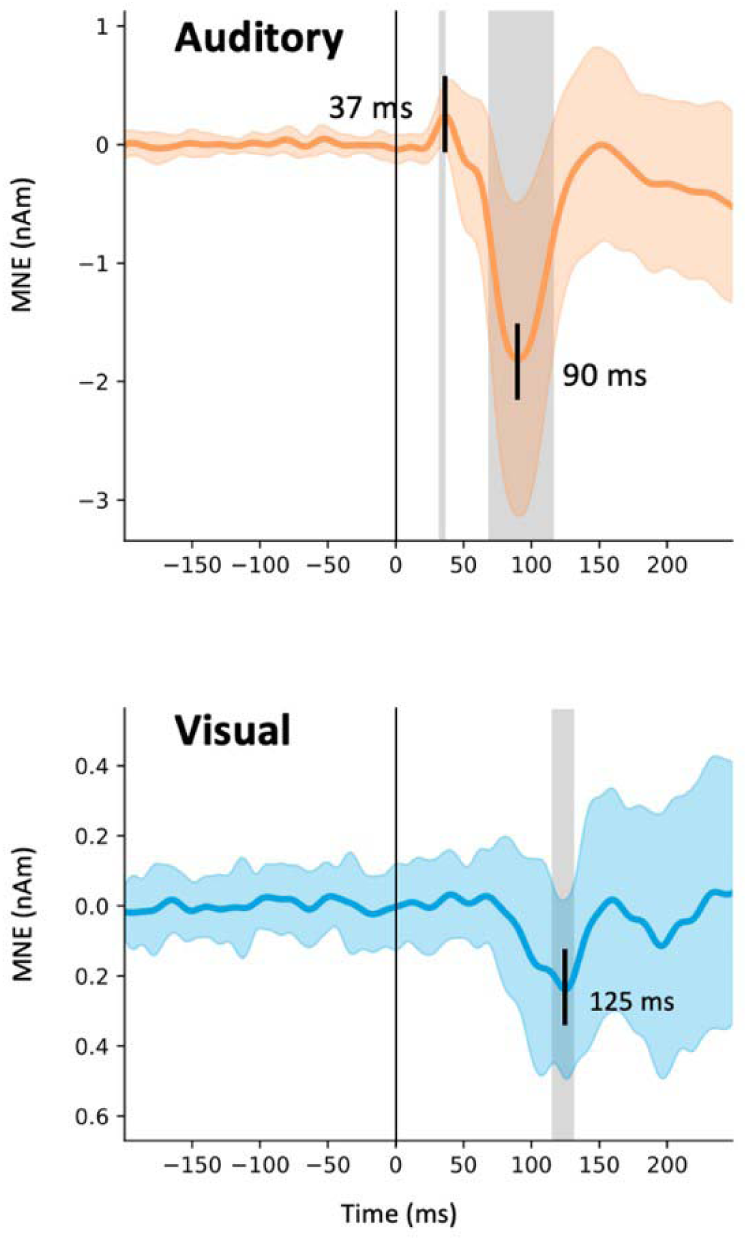
MEG source activity in the auditory cortex. The estimated source waveforms in response to the auditory (orange) and visual (blue) stimuli (mean and standard deviation across subjects, hemispheres, and experiments). Negative values correspond to inward cortical currents, i.e., pointing from the gray matter towards the white matter. The gray shading indicates time points that differing significantly from zero (t-test, p < 0.05, Bonferroni adjusted).

We examined the reproducibility of the estimated source waveforms across the experiments, hemispheres, and individual subjects. MEG source waveforms in the left and the right hemispheres in response to the *Noise/Checkerboard* and *Letter* stimuli are illustrated in **Fig. 3**. The magnitude of the auditory N100m was larger for the *Letter* than for the *Noise* stimuli in the left hemisphere, but similar in the right hemisphere; this lateralization is expected for responses to phonetic vs. non-verbal stimuli (Gootjes et al., 1999; Parviainen et al., 2005). The anatomical overlap of ROIs across subjects (**Fig. 3**, middle panel) suggested that the prominent auditory evoked responses originated mostly in the Heschl’s sulcus and the anterior part of the planum temporale. There were no clear differences in the location of the ROIs between the *Noise/Checkerboard* and *Letter* experiments; however, for the *Letter* stimuli, the location extended to the Heschl’s gyrus in half of the subjects. The peak latencies of the auditory evoked responses were similar within a few milliseconds in both experiments. For the visual evoked response, a negative deflection with the peak latency ranging from 113 to 132 ms was seen in 311 both experiments in both hemispheres.

**Figure 3.**
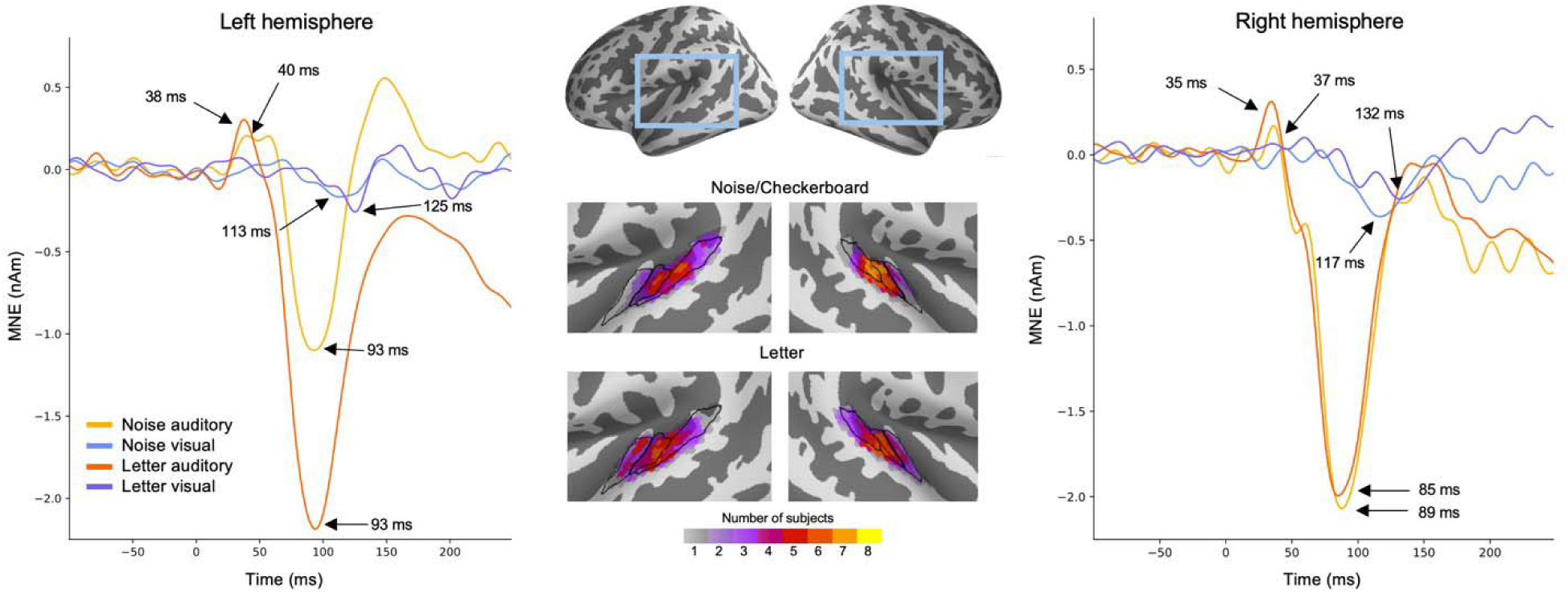
MEG source waveforms in the left and right hemisphere auditory cortex in response to auditory and visual stimulation, shown separately for the Noise/Checkerboard and Letter experiments. The source waveforms were averaged over subjects. The locations of the functional 316 ROIs morphed to common anatomical space (‘fsaverage’ from FreeSurfer) are shown in the middle; the color bar indicates how many subjects’ individual ROIs overlapped at each cortical location. The black lines illustrate the Heschl’s gyrus (anterior), Heschl’s sulcus (middle) and part 319 of planum temporale (posterior).

To evaluate between-subject consistency of the largest defections in the evoked responses in each hemisphere in each experiment, we calculated for each subject the average value over time points within ±10 ms windows around the peak latencies (black dots in **Fig. 4**). The auditory N100m peak was statistically significant in all cases (*Noise*: left hemisphere *p* = 0.027, right *p* = 0.0045; *Letter*: left *p* = 0.027, right *p* = 0.027; *t*-test, False discovery rate (*fdr)* adjusted). For the response to the visual stimuli, the negative peak was statistically significant in the right hemisphere (*Checkerboard*: *p* = 0.040; *Letter*: *p* = 0.027) but not in the left hemisphere (*Checkerboard*: *p* = 0.19; *Letter*: *p* = 0.). The auditory P50m peaks were not significant when calculated separately for the different cases, but they were significant for the grand average responses (see **Fig. 2**).

**Figure 4.**
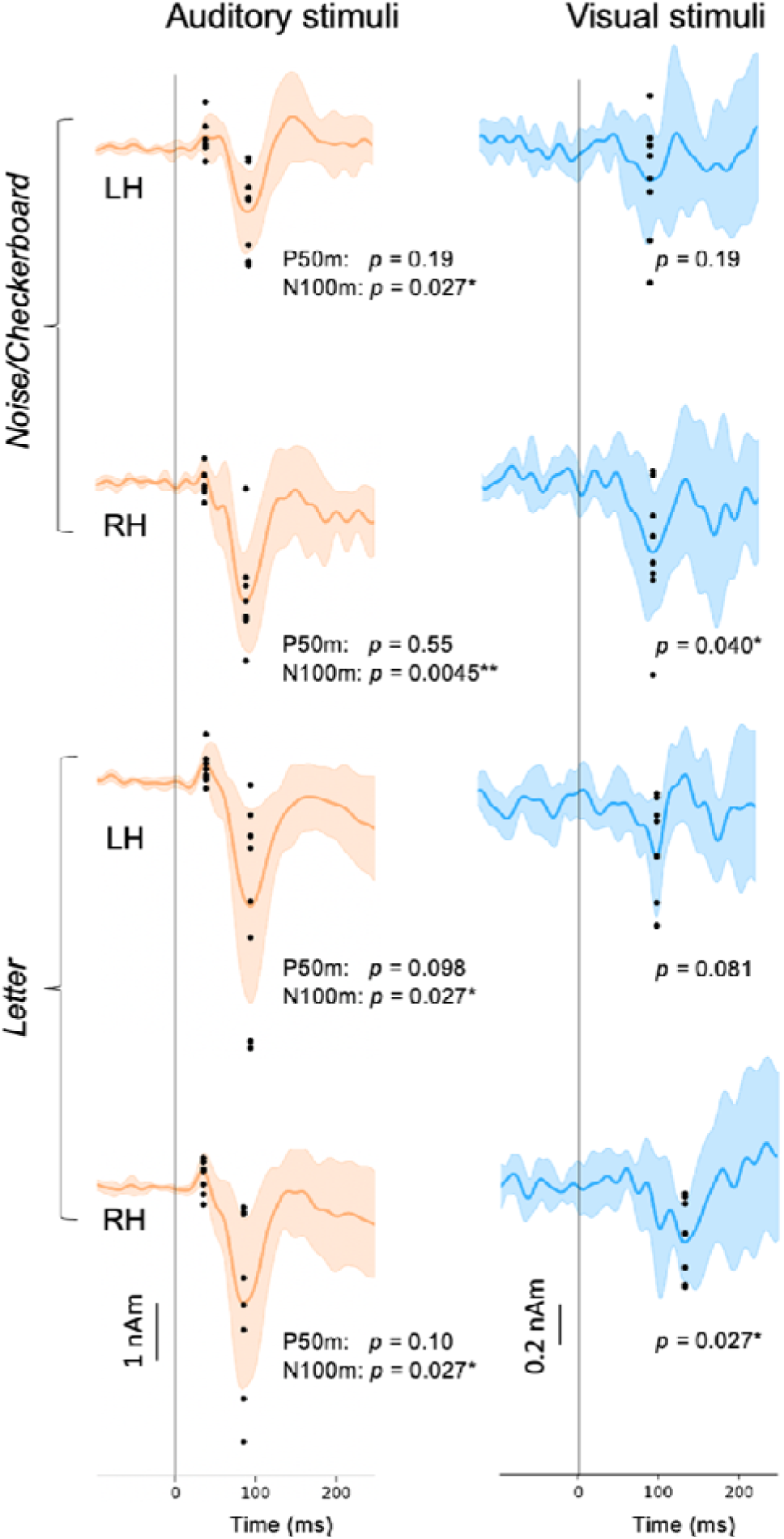
Variation of the estimated source waveforms among individual subjects. The p-values indicate the significance of the response magnitudes at the peak latencies (t-test; fdr adjusted). Continuous lines and shading: mean ± standard deviation across subjects; black dots: response magnitudes for individual subjects, calculated as the average over ±10 ms time windows around the peak latencies. LH: left hemisphere, RH: right hemisphere. **\*** p < 0.05, ** p < 0.01.

The observed weak visual evoked activity in the auditory cortex partially coincided with strong activity in occipital visual cortical regions (**Fig. 5**). The estimated auditory cortex source waveforms could potentially reflect artefactual spread in the MEG source estimates due to activity in other cortical regions responding to the visual stimuli. We examined this possibility in two ways. First, we observed that the time course of the estimated sources for visual cortex ROIs had prominent deflections for both the onset (with peak latencies at ∼100 ms) and the offset (∼400 ms) of the visual stimuli, whereas in the auditory cortex the response was seen mainly for the onset only (**Fig. 5A**). If the onset and offset responses share a common spatial distribution in the occipital cortex, then also the potential artefactual spreading to the auditory cortex is expected to be the similar after the onset and the offset of the visual stimuli. However, this was not found in the data. Second, the spatial maps of the source estimates for the visual evoked responses have a gap between the weak auditory cortex activity and the large occipital cortex activity (**Fig. 5B)**. Artificial spread would be expected to be spatially uniform rather than forming separate foci in the auditory cortex. These observations argue against the possibility of the cross-sensory visual evoked response in the auditory cortex to be artefactually resulting from spread from visual cortex in the source estimates.

**Figure 5.**
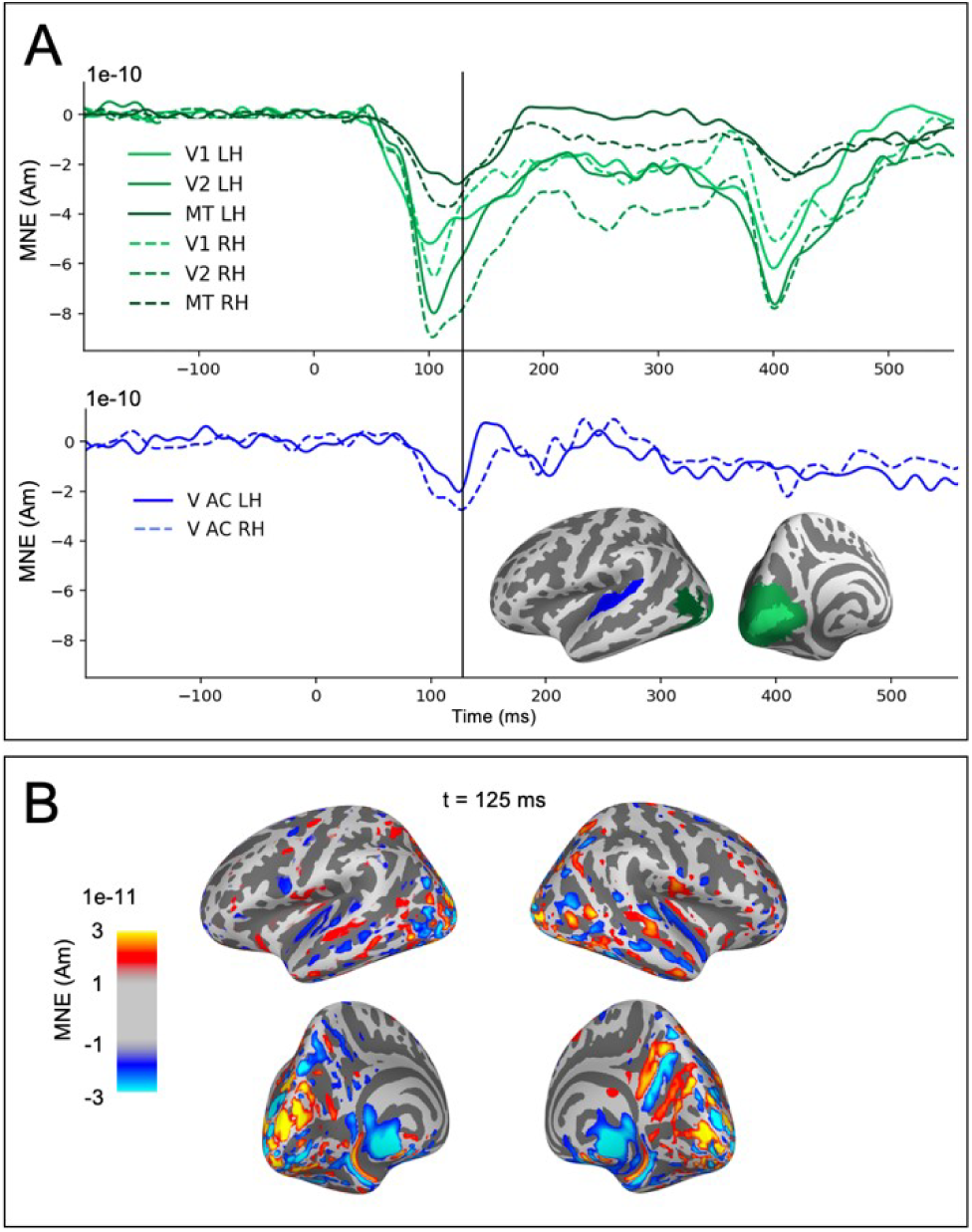
Evaluation of potential artefactual spatial spread in the estimated MEG source activity from visual cortex to the auditory ROIs. (A) Source time-courses (MNE, averaged across subjects and tasks) in response to visual stimuli for occipital areas V1, V2, MT (green) and the auditory cortices (V AC, blue). (B) Spatial maps of the MNE source estimate for the visual evoked activity at the time of the largest peak in the response to visual stimuli in the auditory cortex.

### Neural modeling with HNN

The initial manual tuning values for the mean (and standard deviation) of the time distribution of the inputs were µ_!!_ = 35 (σ_!!_ = 3.0) ms for the FF and µ_!“_ = 75 (σ_!“_ = 13.3) ms for the FB input in the auditory model, and µ_!“_ = 105 (σ_!“_ = 13.3) ms for the FB input in the visual model. The optimal scaling factor was found to be 53 for the auditory and 5 for the visual simulation. Finetuning with Bayesian hyperparameter optimization resulted in only small adjustments to the timing parameters. The optimized values were µ_!!_= 34 (σ_!!_ = 1.0), µ_!“_= 74 (σ_!“_ = 14.0) in the auditory odel, and µ_!“_ = 105 (σ_!“_ = 17.5) in the visual model (**Table 1**). The temporal distributions of 369 puts are depicted in **Fig. 6B**. For both the auditory responses (P50m-N100m) and the visual 370 nses (peaking at 125 ms), the simulated source waveforms captured the main features of 371 the imentally observed MEG results **(Fig. 6A**).

**Figure 6.**
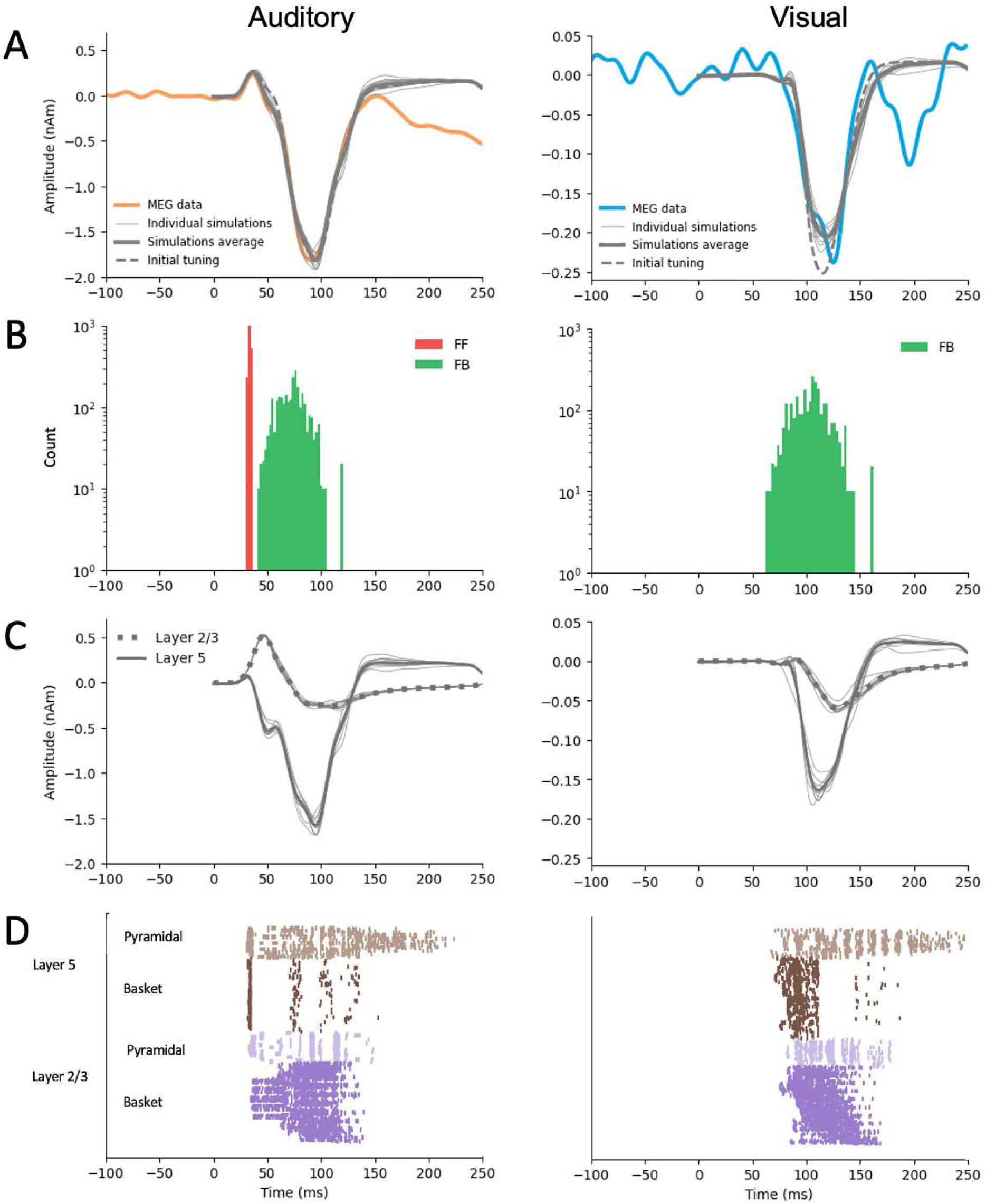
HNN simulations of the auditory cortex activity in response to auditory (left) and visual (right) stimuli. **A:** Simulated source waveforms using the initial manual adjustments to the model parameters (dashed gray lines), after parameter optimization (thick gray: average, thin gray: 10 individual simulation runs), and the measured MEG data averaged over subjects, hemispheres, and experiments (orange: auditory, blue: visual). **B:** Histograms of the timing of the inputs sampled from a Gaussian distribution with a model-specific mean and standard deviation (red: FF, green: FB) **C:** Layer-specific simulations after optimization (green: layer 2/3, purple: layer 5, gray: 10 respective individual simulation runs). **D:** Spiking activity of the pyramidal and basket cells in layers 2/3 and layer 5 (10 simulation runs).

**Table 1.** Comparison of HNN parameters for auditory and visual models. The mean µ and standard deviation σ (milliseconds) describe the temporal distribution of the inputs. Scaling is used to match the simulated dipole to the measured evoked response waveform. E is root mean-square error calculated between simulated and measured waveform. The models are highlighted.

| Model | $\mu_{FF} (\sigma_{FF})$ | $\mu_{FB} (\sigma_{FB})$ | $\mu_{FF2} (\sigma_{FF2})$ | Scaling | RMSE |
| --- | --- | --- | --- | --- | --- |
| A: FF + FB + FF<br>(Kohl et al., 2022) | 47 (3.0) | 81 (13.3) | 151 (11.1) | 1500 |  |
| A: FF + FB + FF | 35 (1.0) | 77 (15.0) | 90 (14.4) | 57 | 0.15 |
| A: FF + FB | 34 (1.0) | 74 (14.0) | - | 53 | 0.072 |
| A: FB | - | 78 (14.8) | - | 38 | 0.23 |
| A: FF | 20 (3) | - | - | 1 | 0.86 |
| V: FF + FB | 42 (1.0) | 99 (12.7) | - | 6 | 0.012 |
| V: FB | - | 105 (17.5) | - | 5 | 0.024 |
| V: FF | 1 (3.0) | - | - | 1 | 0.095 |

Further insights to the generation of the source currents can be obtained by plotting separately the contributions from layer-2/3 and in layer-5 pyramidal cells (**Fig. 6C**) and the sequences of the spiking activity of the four cell types included in the HNN model (**Fig. 6D**). In the model for the auditory evoked response, FF input was assumed to arrive to the auditory cortex through the middle cortical layer and the excite the basal dendrites of the pyramidal cells in both layers 2/3 and 5 **(Fig. 6C**, left). The net result of the FF input was an initial upward (positive) peak. The arrival of the FB input to the distal parts of the apical dendrites of the pyramidal cells resulted in reversal of the net current to be downwards. In the model for the cross-sensory visual evoked response, the FB input arriving distally drove the net source current downwards within the apical dendrites of both layer 2/3 and layer 5 pyramidal cells **(Fig. 6C**, right).

As HNN has a large number of parameters, it is possible that our chosen models are not the only ones that can reproduce the experimentally observed MEG source waveforms. However, HNN can serve us as a valuable hypothesis testing tool to test different models. Alternative models with different combinations of FF and FB inputs are shown in **Fig. 7**, and the corresponding optimized HNN parameters for these are listed in **Table 1**.

**Figure 7.**
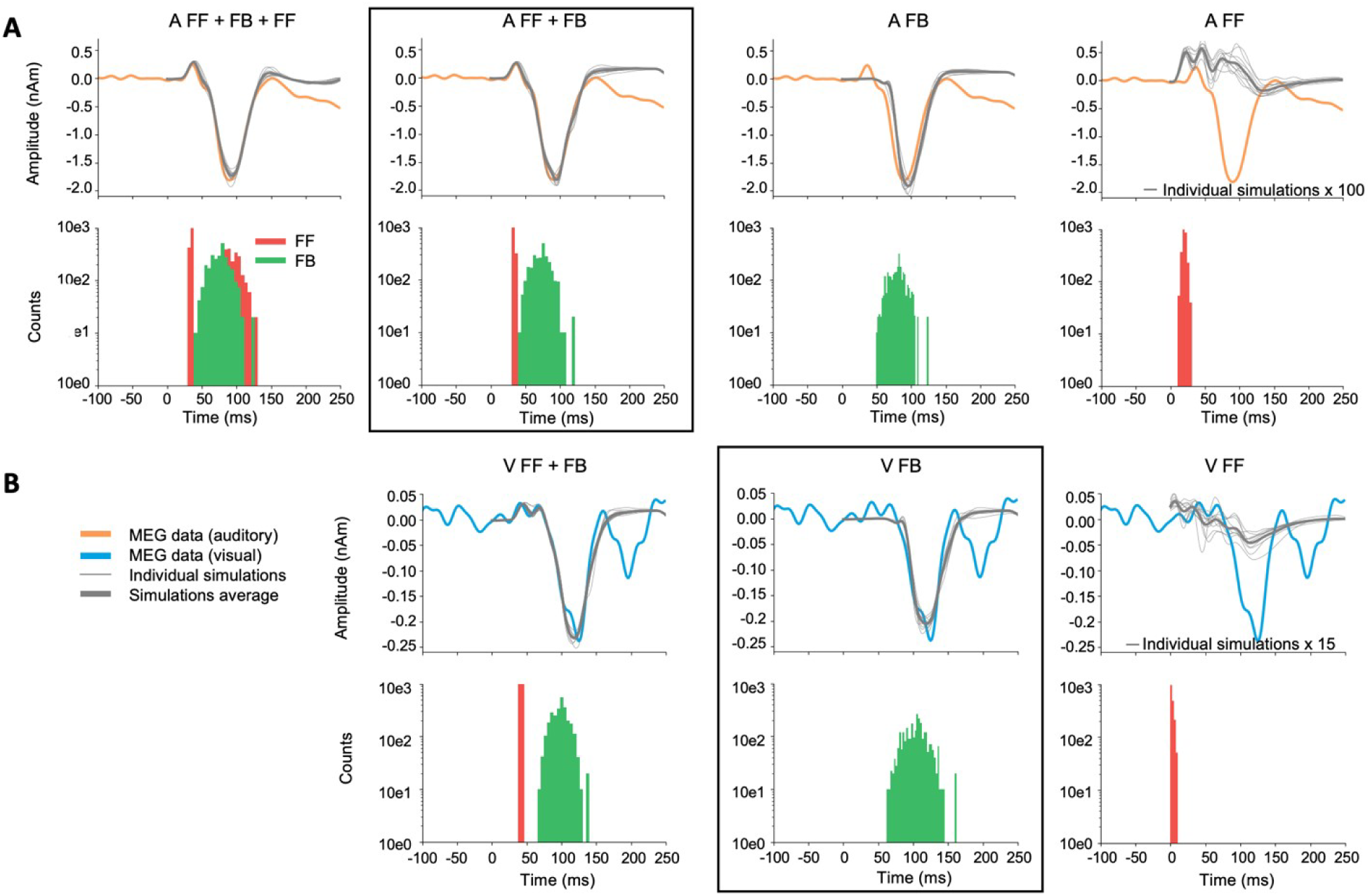
Alternative models for auditory. (**A**) and visual (**B**) responses. The main models (A: FF + FB and V: FB) are framed. The experimentally observed MEG source waveforms (orange: auditory stimulus, blue: visual stimulus) are overlayed with the simulated waveforms (thin gray: 10 individual simulation runs, thick gray: average of the individual runs. Histograms below the waveforms show the temporal distribution of FF (red) and FB (green) inputs to the HNN model of the auditory cortex neural circuit. FF only simulations are scaled to illustrate their waveforms compared with the MEG signal.

For the auditory evoked responses, inclusion of a later second FF input to the model had only little effect on the simulated source waveforms within 0–150 ms (A: FF+FB+FF2 *vs.* A: FF+FB, **Fig. 7A**). Removing the first FF input, however, resulted in a notable difference in the early time window (30–80 ms), during which the first upward deflection was seen in the MEG data. Interestingly, if the FB input was removed, the FF input alone could not produce response waveforms similar to those observed empirically. As the optimal scaling factor for the FF only model was 1, **Fig. 7** (right column) shows the model scaled up in order to illustrate how the waveform looks like compared with the MEG response. Thus, the FB input seems to have an essential role in the generation of the evoked responses studied here.

For the visual evoked response, the difference between models with and without an FF input (V: FF+FB *vs.* V: FB) was most pronounced in the early part (30–80 ms) of the simulated source waveforms **(Fig. 7B**). However, although the V: FF+FB model slightly improved the fit to the measured MEG signal in comparison with V: FB, considering the magnitude of the response with the baseline noise level (see **Fig. 2**) suggests that the additional FF input in the model for the response to the visual stimuli may be mostly explaining just noise in the data. Using a nonparametric resampling approach, a significant difference between FF+FB *vs.* FB was found for the auditory models (*p* < 0.001) but not for the visual models (*p* = 0.39). In other words, early FF input did not significantly improve the model fit to the response to visual stimuli. Thus, these results support our main hypothesis that the response to the auditory stimuli results from a combination of FF and FB inputs to the auditory cortex, whereas the cross-sensory visual response can be explained with just FB input to the auditory cortex.

## Discussion

The MEG data revealed a cross-sensory event-related response in the auditory cortex, peaking at about 125 ms after the appearance of the visual stimuli. The direction of the estimated source current for this response was the same as for the auditory N100m response, pointing from the cortical gray matter towards the white matter. The main shape of the visual evoked response waveform could be reproduced by an HNN model with FB-type input, whereas for the biphasic P50m-N100m auditory evoked response both FF and FB inputs were needed. The experimental and modeling results are consistent with the hypothesis that cross-sensory visual input to the auditory cortex is of FB type (Schroeder and Foxe, 2002).

### Characterization of cross-sensory visual evoked activation in auditory cortex

Recently, Kohl et al. presented an HNN model with a sequence of FF and FB inputs explaining several properties of auditory evoked responses in the auditory cortex (Kohl et al., 2022). With only minor adjustments to the input timings and the overall scaling, the model could be adapted to explain the MEG source waveforms for the auditory evoked responses observed in the present study. A sequence of FF-FB (and –FF) inputs has been shown to model well also somatosensory responses in the somatosensory cortex (Jones et al., 2007). In contrast, to explain the early part of the cross-sensory visual response in the auditory cortex, we found that a model with only an FB input, without a preceding FF input, was adequate. The FB-type characteristics is consistent with previous NHP electrophysiological studies (Schroeder and Foxe, 2002). Multi-contact electrode recordings in the macaque have shown early activity in the granular (middle) layer of auditory cortex in response to auditory stimuli, suggesting FF-type input, whereas cross-sensory visual evoked activity appeared first in supra– and infragranular layers (Schroeder and Foxe, 2002). Similar laminar properties in the auditory cortex have also been seen in human fMRI studies (Gau et al., 2020; Chai et al., 2021; Lankinen et al., 2022). In the high-field laminar fMRI study of Lankinen et al. (2022), which used the same stimuli as in the *Noise/Checkerboard* experiment in the present MEG study, BOLD signal depth profiles in the auditory cortex showed different curvature for auditory vs. visual stimuli, consistent with the hypothesized difference in the FF vs. FB type inputs.

There are several possible neural pathways for the visual evoked activity to reach the auditory cortex. The relatively long latency of the visual response observed here is consistent with what would be expected from input from higher-order polysensory areas such as the superior temporal sulcus (Foxe and Schroeder, 2005). However, the present analyses focusing on activity within auditory cortex only do not reveal the origin of the inputs to the auditory cortex. That type of information could be deduced, e.g., from Granger-causality measures between estimated source waveforms in multiple cortical areas (Milde et al., 2011; Gow and Nied, 2014; Michalareas et al., 2016).

Interestingly, NHP studies have shown different characteristics for visual and somatosensory cross-sensory inputs to the auditory cortex: FB-type for visual but FF-type for somatosensory (Schroeder and Foxe, 2002). The role of different types of cross-sensory inputs to the auditory cortex may have important implications to theories of multisensory processing (Schroeder and Foxe, 2005). There appear to be multiple ways how cross-sensory processes may be influenced by the hierarchical organization among brain areas. FB-type inputs are commonly associated with modulatory influences, whereas FF-type inputs are more directly related to sensory information (Schroeder and Foxe, 2005).

### Complementary approaches to noninvasive detection of FF and FB processes

The present approach of combining MEG and cellular-level computational modeling complements other non-invasive methods for studying the organization of cortical processes in the human brain. The millisecond-scale time resolution of MEG and EEG enables the investigation of fast dynamics of the brain activity, which is not attainable with hemodynamic fMRI. High-field fMRI, however, can provide laminar-level spatial resolution for making inferences about FF and FB activity (see e.g., De Martino et al., 2018; Lawrence et al., 2019b; Norris and Polimeni, 2019). With certain strong assumptions about the location and extent of the spatial distribution, layer-specific source localization in MEG has also been demonstrated (Bonaiuto et al., 2018a; Bonaiuto et al., 2018b). FF/FB influences can also be inferred from directed connectivity measures for MEG source estimates at specific frequency bands (Michalareas et al., 2016).

The present results also support the view that the direction of MEG source waveforms can be useful for inferring information about the hierarchical organization of cortical processing (Ahlfors et al., 2015). In particular, FF-type input to the supragranular layer, with excitatory synaptic connections to the distal part of the apical dendrites of pyramidal cells, is likely to be a major contributor to the downward-directed MEG source currents (Lopes da Silva, 2010; Ahlfors and Wreh, 2015). There was a general correspondence between the source direction and the type of input in the HNN model: the outward directed source current during the auditory P50m response was associated with FF input in HNN, whereas FB inputs were needed to model the inward source currents during the auditory N100m and the visual response peaking at 125 ms. A close relationship between the direction of MEG source currents and FF– vs. FB-type inputs has also been found in HNN modeling of somatosensory response in the primary somatosensory cortex (Jones et al., 2007). Furthermore, the direction of the MEG source currents in inferior occipitotemporal cortex has been found to reverse between two experimental conditions for which a cognitive neuroscience theory for visual object recognition predicted FF vs. FB inputs to the area (Ahlfors et al., 2015).

### Limitations of the current study

Localizing weak cross-sensory visual evoked activity in the auditory cortex is challenging because of potential interference in the MEG source estimate from the partially coinciding occipital cortex activity. However, both the shape of the time courses and the patterns in the spatial distributions of the source estimates (see **Fig. 6**) suggested that it was unlikely that the visual evoked activity in the auditory cortex was due to artefactual long-range crosstalk caused by spatial spread in the source estimates. Short-range spread in the source estimates can also confound the interpretation of the source waveforms. If the true location of the visual responses were not within the auditory cortex ROI, but, e.g., in the opposite side of the superior temporal gyrus, the source direction could become incorrectly identified. Combining MEG with high-resolution fMRI could help to confirm the location of the activity. It is also possible that there was simultaneous activity in multiple auditory areas in the supratemporal plane. Most of the individual subjects’ ROIs were located directly at the primary auditory regions, at or near the at Heschl’s sulcus, being thus slightly different than the auditory association area just posterior to primary auditory region studied by Schroeder and Foxe (2002). However, it has been shown in monkeys that such FF type patterns are typical throughout the core and belt regions of auditory cortex (Schroeder et al. 2001). Without further data, e.g., intracranial recordings, it is difficult to conclusively resolve the locations of the sources of the observed cross-sensory MEG response.

HNN, and biophysical computational neural modeling in general, has two challenges of opposite nature: the neural circuit model is complex, with a large number of adjustable parameters, and yet the model is a simplified representation of the cortical circuitry. We used neural circuit parameters of the pre-tuned model for auditory evoked responses in the auditory cortex by Kohl et al. (2022) and only adjusted a small number of selected parameters, focusing on the timing of the FF and FB inputs. Given the limited SNR of the experimental source waveforms, we did not attempt to vary the neural connectivity parameters. We cannot exclude the possibility that there could be some combinations within the high-dimensional parameter space that could explain the responses with a very different circuit model than the one reported here. Useful in future studies, it has been recently demonstrated that combining simulation-based inference (SBI) to HNN modeling can help in parameter estimation (Tolley et al., 2023).

We modeled only one local region (auditory cortex) receiving one-directional external inputs. To determine where the inputs are arriving from and where the information will be sent, directional connectivity analyses between multiple regions would be needed. Thus, further studies would be necessary to connect other areas of interest to the network. Furthermore, combining MEG with layer-specific fMRI could provide complementary information which could help to build a more detailed picture of the FF/FB influences.

## Conclusions

The combined MEG and HNN modeling results support the hypothesis that cross-sensory visual input to the auditory cortex is of FB type. The results also illustrate how the dynamic patterns of the estimated MEG/EEG source activity can provide information about the characteristics of the input into the cortical areas in terms of hierarchical organization among the cortical areas. Avenues for future research could include connecting other areas of interest to the network, calculating directed (effective) connectivity measures between cortical areas specifically, and combining complementary information from MEG data with layer-specific fMRI to build a more detailed picture of the FF/FB influences.

## Acknowledgments

Supported by R01DC016765, R01DC016915, R01DC017991, R01NS126337, P41EB030006, P41EB015896, S10OD030469. We thank Dr. Stephanie Jones for useful discussions.

